# PyVADesign: a Python-based cloning tool for one-step generation of large mutant libraries

**DOI:** 10.1101/2025.04.04.647202

**Authors:** R.C.M. Kuin, M.H. Lamers, G.J.P. van Westen

## Abstract

**Motivation:** The generation and analysis of diverse mutants of a protein is a powerful tool for understanding protein function. However, generating such mutants can be time-consuming, while the commercial option of buying a series of mutant plasmids can be expensive. In contrast, the insertion of a synthesized double stranded DNA (dsDNA) fragment into a plasmid is a fast and low-cost method to generate a large library of mutants with one or more point mutations, insertions, or deletions.

**Results:** To aid in the design of these DNA fragments we have developed PyVADesign: a Python package that makes the design and ordering of dsDNA fragments straight-forward and cost-effective. In PyVADesign, the mutations of interest are clustered in different cloning groups for efficient exchange into the target plasmid. Additionally, primers that prepare the target plasmid for insertion of the dsDNA fragment, as well as primers for sequencing, are automatically designed within the same program.

**Availability:** PyVADesign is open source and available at https://github.com/CDDLeiden/PyVADesign/.

## 1 Introduction

Proteins are essential to all biological systems due to their diverse functional roles, ranging from catalyzing biochemical reactions to regulating cellular processes and enabling communication between cells (Morris *et al*., 2022; Agarwal, 2006; Westermarck *et al*., 2013). Mutations in proteins can lead to altered functions, which can be detrimental or in some cases provide evolutionary advantages, thus playing a crucial role in both disease mechanisms and adaptive evolution (Loewe and Hill, 2010). Consequently, studying the behavior of mutants provides valuable insights into protein function, aids in protein engineering, helps explain disease development, and elucidates drug resistance (Sellés Vidal *et al*., 2023; Sen *et al*., 2022; Tait, 1999). Although generating mutants can be time-consuming, it remains a fundamental aspect of molecular biology (Forloni *et al*., 2019).

Mutations are traditionally created through site-directed mutagenesis by PCR (Weiner et al., 1994; Bachman, 2013). As all mutants are introduced in a per-case basis, creating different mutations can become time-consuming as each mutations demands a unique set of primers and PCR reaction, which becomes labour-intensive for larger sets of mutants. In addition, the generation of multiple mutations in a gene using a single PCR is limited to residues that are close to each other in the sequence. For multiple mutations that are spaced further apart, there is no other options than to go through multiple rounds of mutagenesis. In contrast, synthesis of whole genes or whole plasmid offers a fast alternative but can become expensive when multiple mutant versions are required.

Alternatively, synthesis of short double stranded DNA (dsDNA) fragments ranging from a few hundred to thousands of base pairs (bp), combined with easy-to-use cloning methods such as in-vivo assembly (IVA) (García-Nafría *et al*., 2016) offers a quick and economic approach to generate large mutant libraries with one or more point mutations, as well as insertions or deletions (Hughes and Ellington, 2017). To streamline this process, we created PyVADesign; a Python package that makes design and ordering of dsDNA fragments straightforward. Using PyVADesign, mutations of interest are clustered within the target gene and a dsDNA fragment is designed for each variant, see Figure 1. The program also automatically generates primers to prepare the expression plasmid for dsDNA fragment insertion, as well as primers for validation by DNA sequencing. the tool is open-source and user-friendly for scientists of all backgrounds.

**Figure 1.**
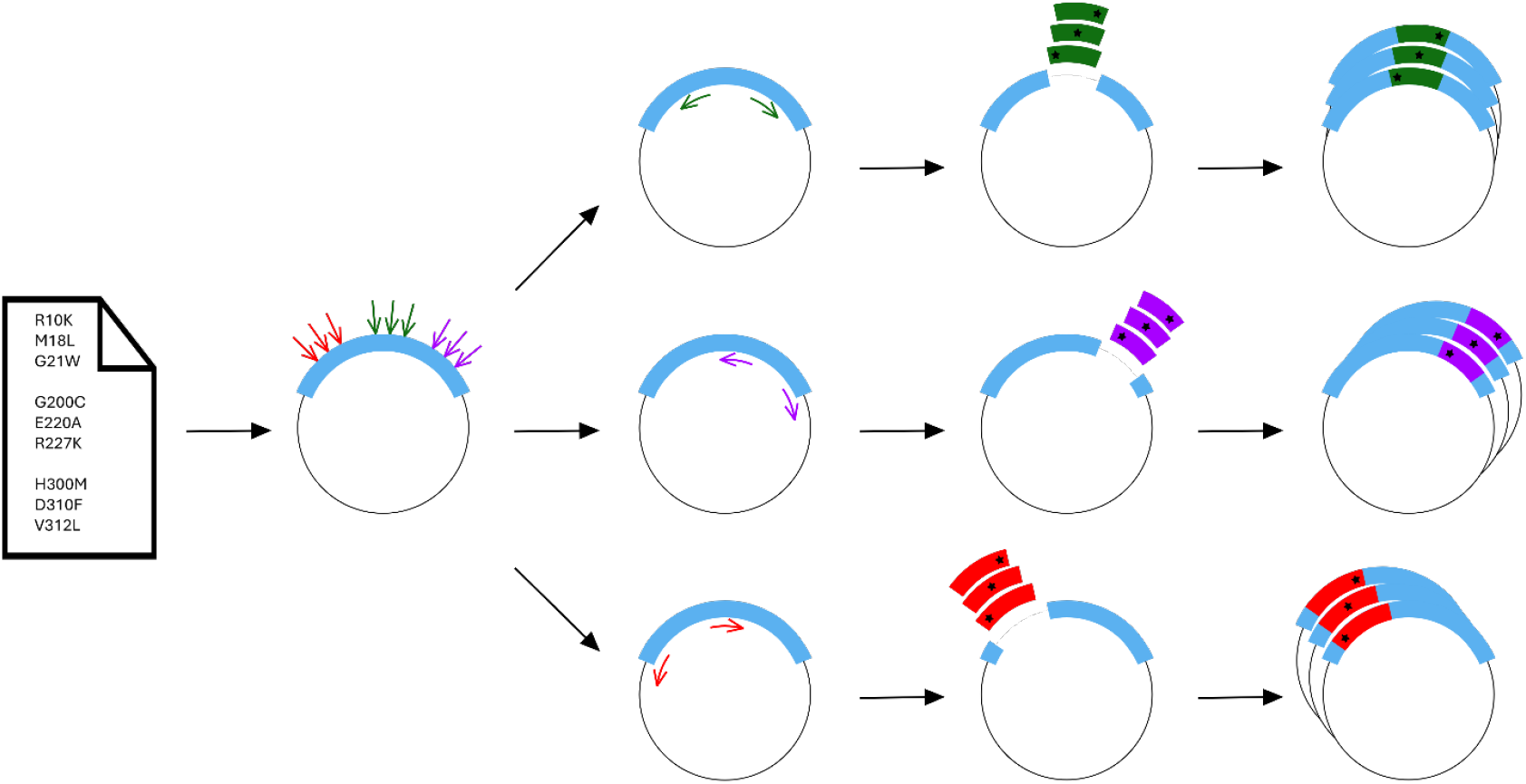
Graphical representation of the PyVADesign workflow. User’s input includes a list of desired mutations and a plasmid sequence with gene of interest (blue). PyVADesign clusters the mutations on the gene of interest without explicit user input. Next, the dsDNA fragments and primers are designed to prepare the plasmid for dsDNA fragment insertion (shown in green, purple and red colours). To linearize the input plasmid for insertion of the dsDNA fragment, PCR reactions are performed using the PyVADesign primers. Finally, the dsDNA fragment can be inserted in the linearized plasmid in a single step.

## 2 Methods

### 2.1. dsDNA fragment design process

First, the desired mutations are grouped together into clusters using K-medoids clustering (Kaufman and Rousseau, 1990) in scikit-learn-extra (The scikit-learn-extra development team, 2020). Starting with two clusters, we assess if the resulting cluster sizes meet the size requirements. This process is repeated with an increasing number of clusters, until the cluster sizes become smaller than the minimum size of the dsDNA fragment. Using this approach, fragment regions are created that cover a maximum of mutations and leave out regions for which no mutations are designed (Supplementary Fig. 1). For two or more simultaneous mutations, constraints are generated for paired data points, modifying the distance metric by reducing the distance between them to promote their co-clustering. The cluster selection depends on user preference and can be based on either fragment-quantity or fragmentlength. The quantity-optimization option selects a clustering based on the number of different fragment regions, aiming to reduce the number of required PCRs. The length-optimized option prioritizes minimizing the total number of base pairs per region, which results in a higher number of smaller fragment regions compared to the quantity option and which are cheaper than the larger fragments.

After clustering, the mutations are assigned to a fragment region and a dsDNA fragment is created for each mutant. For each mutation, the most frequently used codon in the selected (expression) organism is determined by retrieving the genome from the NCBI Entrez database (Sayers *et al*., 2021) and calculating the relative frequencies using Biotite (Kunzmann and Hamacher, 2018). Flanking sequences are added to both ends of the dsDNA fragment that are required for the insertion into the target plasmid via IVA cloning (García-Nafría *et al*., 2016). Lastly, two silent mutations are included on both ends of the dsDNA fragment as an easy identification tool, bypassing the need to sequence the entire insert.

### 2.2. Primer design

After the dsDNA fragment design, two sets of primers are designed: one set of primers to open-up the target plasmid to allow for insertion of the dsDNA fragment and one set of primers for sequencing of the created plasmid. Primers are designed using the Primer3 package (Untergasser *et al*., 2012; Rozen and Skaletsky, 2000). Settings for both primer sets can be adapted to user preferences through a settings file, allowing for customization of parameters such as optimal primer length, melting temperature, and GC content. Within the Primer3 package, the primers are also checked for the formation of secondary structure and off-target binding sites.

### 2.3. DNA synthesis

All dsDNA fragments and primers were ordered from IDT^TM^ (Leuven, Belgium). The dsDNA fragments were delivered in a 96-well plate at 10 ng/µL in nuclease-free water.

### 2.4. Preparation of target plasmid

The target plasmid was opened up using the designed primer set through PCR. PCR was performed by using KOD Hot Start Master Mix (Sigma-Aldrich), according to the manufacturer’s instructions. Briefly, each reaction contained 10 ng plasmid, 0.3 μM of each primer, 25 μL of KOD Hot Start Master Mix in a total volume reaction volume of 50 μL. PCR consisted of 20 cycles denaturation at 95 °C, annealing at 65 °C and extension at 72 °C, with a final extension step of 225 seconds at 72 °C. Reaction mixtures were treated with DpnI for 60 minutes to remove the parent plasmid. The linear PCR fragments were purified from a 1% agarose gel using the Wizard® SV Gel and PCR Clean-Up System (Promega) and stored at −20 °C until use.

### 2.5. Assembly of mutant plasmid

To assemble the mutant plasmid, 20 ng of the synthetic dsDNA fragment and 100 ng opened up target plasmid were combined and transformed into *E. coli* DH5α cells using a heat shock method (Froger and Hall, 2007). Single colonies were picked and grown in 5 ml Luria-Bertani (LB) at 37 °C overnight. DNA was extracted using the QIAprep Spin Miniprep Kit (Qiagen). All plasmids were sequenced by Illumina sequencing.

## 3 Results

The software package is written in Python version 3.9 and is organized into modules, each handling a separate task. If no customization is required, the package offers a command line interface for quick setup of experiments with default settings so the package is easy to use for molecular biologists. For more complex experimental designs, a settings file can be supplied to customize both the dsDNA fragment design process and primer parameters. Both the dsDNA design process and primer design process are also described in a step-by-step tutorial in the GitHub repository. The tool has been tested on Windows and Unix-based systems to ensure cross-platform functionality.

### 3.1. User input and considerations

To successfully run the design process, a text file with the mutations of interest is required listing various types of mutations, including single point mutations, multiple point mutations, insertions, and deletions, see the Supporting Data for an example file. It is important to note that, depending on the sizes of the dsDNA fragments, paired mutations cannot be too far apart as they need to fall within the same cluster to be incorporated into a single dsDNA fragment. The gene of interest must be provided in FASTA format, and the plasmid sequence containing the gene of interest should be available in GenBank (.gb) or SnapGene (.dna) format. Values for dsDNA fragment lengths and price per base pair can be adjusted to match the specifications of your DNA supplier.

### 3.2. Output

For every mutant a complete vector sequence file in GenBank and SnapGene format is generated that can readily be imported into a sequence manager such as Benchling (The Benchling development team, 2024) or SnapGene (The SnapGene development team, 2024). These vector sequences contain the mutation of interest as well as the cloning and sequencing primers. For ordering of the dsDNA fragments and primers a complete list of sequences is generated to streamline the ordering process. The localization of the fragment regions as well as the mutations within the gene of interest is also visualized using the plotting library DnaFeaturesViewer to easily interpret the results (Zulkower and Rosser, 2020).

### 3.3. Validation with a random synthetic test set

To test PyVADesign and to estimate the limits of the tool regarding the maximum insert and deletion length and the distance between paired mutations, we generated a synthetic dataset. For this, we introduced five random point mutations along with an additional insertion, deletion, or paired mutation of a defined length. Specifically, we systematically increased the length of the insertion, deletion, or paired mutation from 1 to 600 in steps of 5, repeating the process for each length. This procedure was performed for three different genes and repeated 10 times. Next, we designed dsDNA fragment regions for each mutated plasmid using the default settings and calculated the percentage of cases where dsDNA fragments could be designed, see Supplementary Table 1. We found that dsDNA fragments could be designed for deletions up to 185 residues, inserts of 115 residues and paired mutations within 230 residues with a success rate over 90 %. Starting from a list of input mutations, the design of the dsDNA fragments and primer sets for opening-up the plasmids and sequence validation takes approximately 2 minutes.

### 3.4. Experimental validation

To showcase the applicability of our approach, we designed 24 mutations in the DNA polymerase DnaE1, an essential protein of *Mycobacterium smegmatis*. The PyVADesign parameters, the fragment regions, dsDNA fragments and associated primers can be found in Supplementary Fig. 1, and Supplementary Table 2-3.The 24 mutations contained 18 point mutations, four double mutations and two deletions. The 24 mutations were distributed over three fragment regions. The dsDNA fragments were combined with its appropriate target plasmid and transformed into DH5α cells. Using this approach, all 24 mutant plasmids were generated within a span of two weeks.

## 4 Conclusion

In this paper, we present a novel approach for generation of large mutant libraries using commercially available dsDNA fragments in combination with an IVA cloning approach. We implemented a Python tool that automates the process of designing dsDNA fragments and associated primers. Size limits of the dsDNA fragments for inserts and deletions were evaluated, and the approach was experimentally validated. In some cases, PyVADesign fails to identify a clustering solution for the input mutations due to the distances between them. To address this, and to surpass the current size limits, the design process could be adapted to allow for multifragment assembly.

As an open-source tool with minimal user input requirements, PyVADesign is intended to be accessible to a diverse set of users. Additionally, by reducing the number of required PCRs, the tool enhances both the efficiency and scalability of mutant generation.

## Acknowledgements

The authors thank Adam J. Fountain for his suggestion of using eBlocks for rapid creation of gene variants. The authors also thank Roelof van der Kleij for his help using the university IT infrastructure.

