## Supplementary Data for "PyVADesign: a Python-based cloning tool for one-step generation of large mutant libraries"

**Table 1.** Success rates on synthetic dataset. Success rates (50%, 75%, 90%) of designing dsDNA fragments on a synthetic dataset. For each length (with steps of five) we used 30 different random input mutations to estimate the limits of PyVADesign using the default parameters.

| | Success >50% | Success >75% | Success >90% |
|---|---|---|---|
| Insertion length (bp) | 220 | 170 | 115 |
| Deletion length (bp) | 350 | 245 | 185 |
| Distance between paired mutations (bp) | 350 | 315 | 230 |

**Table 2**. Primers used to linearize the expression plasmid and to validate dsDNA fragment insertion.

| Primer name | Direction | Primer set | Sequence (5' to 3') |
|---|---|---|---|
| fw_DNABlock-1 | Forward | Opening-up target plasmid | GGTGACGGCTACTACCT |
| rv_DNABlock-1 | Reverse | Opening-up target plasmid | CTCCACGCCGATGATCG |
| fw_DNABlock-2 | Forward | Opening-up target plasmid | CGCGCTGTACCGGCC |
| rv_DNABlock-2 | Reverse | Opening-up target plasmid | TCGATATCGATGTCGGGCAT |
| fw_DNABlock-3 | Forward | Opening-up target plasmid | CGATCTCTTCGGCAGCAATG |
| rv_DNABlock-3 | Reverse | Opening-up target plasmid | GCGGGATAGTTGGCCTTGAG |
| seq_fw_DNABlock-1 | Forward | Sequence validation | ACTGGCCTTGTGTTAAAAATGG |
| seq_rv_DNABlock-1 | Forward | Sequence validation | CACTTGTCGGCTGCGTA |
| seq_fw_DNABlock-2 | Forward | Sequence validation | TGACGGTCTGCTCGAGA |
| seq_rv_DNABlock-2 | Forward | Sequence validation | ATGTCTTGGCCGACCGA |
| seq_fw_DNABlock-3 | Forward | Sequence validation | ATGGACTTCCTGGGCCT |
| seq_rv_DNABlock-3 | Forward | Sequence validation | GCGCACTCACCTTGTCA |

**Table 3**. PyVADesign parameters used for the generation of dsDNA fragments for experimental validation

| Parameter name | Description | Value |
|---|---|---|
| max_DNABlock_length | Maximum dsDNA fragment length | 1500 |
| min_DNABlock_length | Minimum dsDNA fragment length | 300 |
| min_overlap | Minimum overlap between dsDNA fragment and plasmid | 25 |
| amount_optimization | Selecting the lowest number of fragment regions | True |
| silent_mutations | Introducing silent mutations at the beginning and end of dsDNA fragment | False |

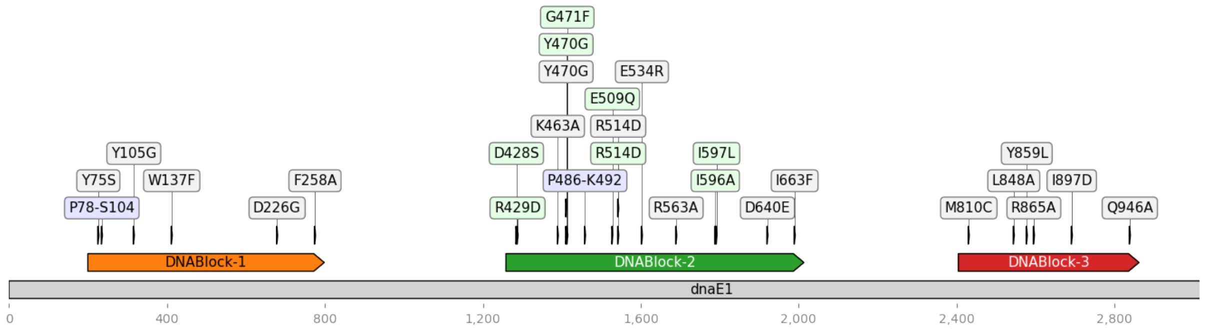


**Figure 1**. Overview of the gene fragments corresponding to the selected mutations used for experimental validation. dsDNA fragments are visualized using DnaFeaturesViewer (Zulkower and Rosser 2020). The target gene is shown in grey, with amino acid numbering indicated. Mutations are represented as follows: single point mutations in grey, paired point mutations in green, and inserts in blue.
